# Music predictability and liking enhance pupil dilation and promote motor learning in non-musicians

**DOI:** 10.1101/812834

**Authors:** R. Bianco, B.P. Gold, A.P. Johnson, V.B. Penhune

**Author notes:** Correspondence: Roberta Bianco.

## Abstract

Humans can anticipate music and derive pleasure from it. Expectations facilitate movements associated with anticipated events, and they are linked with reward, which may also facilitate learning of the anticipated rewarding events. The present study investigates the synergistic effects of predictability and hedonic responses to music on arousal and motor-learning in a naïve population. Novel melodies were manipulated in their overall predictability (predictable/unpredictable) as objectively defined by a model of music expectation, and ranked as high/medium/low liked based on participants’ self-reports collected during an initial listening session. During this session, we also recorded ocular pupil size as an implicit measure of listeners’ arousal. During the following motor task, participants learned to play target notes of the melodies on a keyboard (notes were of similar motor and musical complexity across melodies). Pupil dilation was greater for liked melodies, particularly when predictable. Motor performance was facilitated in predictable more than unpredictable melodies, but liked melodies were learned even in the unpredictable condition. Low-liked melodies also showed learning but mostly in participants with higher scores of task perceived competence. Taken together, these results suggest that effects of predictability on learning can be overshadowed by effects of stimulus liking or task-related intrinsic motivation.

## 1 Introduction

Through passive exposure to music, we implicitly develop models about its structure ^1,2^. These models allow both listeners to generate expectations about upcoming musical events ^3–5^, and trained musicians to better plan and learn musical actions ^6–9^. Expectations are also linked to the experience of musical pleasure ^10–12^, as neuroimaging evidence shows that musically evoked pleasure relies on the cross-talk between neural systems responsible for prediction with those responsible for reward ^13^. Importantly, rewarding stimuli increase arousal ^14^, and are also better learned ^15^. Given the inherent link between predictability and pleasure in music, here we aim to assess their contributions to learning with the hypothesis that motor learning in naïve population may benefit from the implicit musical expectations and hedonic responses derived from music.

A large body of theoretical and experimental work suggests that agents use internal psychological models to make sense of perceptual inputs and respond to them. Through implicit statistical learning, the brain continuously scans the environment for regularities and acquires probabilistic models of the world without deliberate effort or awareness ^16–18^. For example, based on these internal models, the brain can compute the statistical distribution of sequential phenomena ^19,20^ enabling it to predict unfolding sensory events, thereby reducing its uncertainty ^21^. In particular, according to predictive coding theory ^22^, by comparing top-down predictions with the actual sensory input, internal predictive models contribute to the selection of relevant bottom-up inputs ^23^, and error signals are used to update predictions, construct new models, and guide subsequent actions. Internal models can aid motor planning ^24,25^ and it has even been shown that visual statistical learning can inform the motor system to better predict upcoming actions ^26,27^.

Similar cognitive mechanisms are likely to play a role in musical perception and performance. When listening to music, humans entertain a number of predictions, or hypotheses, about future musical events ^1,28^, and these predictions are subject to refinement and learning on different time scales and at different levels of sophistication. For example, musical predictions vary as a function of the musical culture one is exposed to ^29–31^ and can adapt to novel musical styles ^32^. Furthermore, musical predictions are optimized by expertise ^33^, and can even vary across the lifespan ^34^. With regard to performance, it has been shown that musical regularities facilitate movement selection ^35,36^. This facilitation is supported by associations between movement and ensuing effects formed through coupling of motor and sensory cortices ^36–39^. Moreover, studies in trained musicians show that internal predictive models allow long-range motor planning of entire musical sequences ^6,7,40–43^. Thus, motor performance in experts appears to be guided by predictions based on learned internal models of music, suggesting that also initial stages of learning may benefit from them.

Expectations are also linked to the experience of musical pleasure ^10–12,44^. Pleasurable music strikes a balance between predictable events, which allow listeners to form expectations, and moderately unpredictable events that produce surprise. For example, a single repeating note is very predictable, but may not be very enjoyable, and similarly, transitions to unrelated notes may be perceived as unpleasant ^45,46^. Indeed, musical pleasure seems to vary with stimulus complexity – e.g., harmonic or rhythmical predictability – as an inverted-U function, with maximum liking occurring at intermediate levels of complexity ^47–50^. Neural evidence suggests that musical surprises induces liking by engaging the reward system ^51^, via distinct phases of dopamine transmission during the anticipation and enjoyment of listeners’ favorite musical moments ^52^. Connectivity analyses suggest that it does so in cooperation with fronto-auditory systems responsible for predictions during listening ^13^. The role of dopamine, which has been widely linked to the ‘wanting’ and ‘learning’ components of reward ^53^, has been also directly linked with abstract hedonic responses to music (i.e., liking) ^54^. Furthermore, a recent study showed that pleasant musical moments activate the noradrenergic arousal system, as revealed by increased pupil dilation during passive listening ^14^. A concomitant increase of arousal and reward systems during pleasant music seems plausible given the known anatomical and functional link between dopaminergic and noradrenergic subcortical nuclei ^55,56^, and may explain findings of better memory for rewarding than neutral musical excerpts ^57^. Although rarely used to measure response to long stimuli – such as a melody ^58,59^, pupil dilation may thus be powerful to continuously track hedonic responses to unfolding music ^60^.

These evidence that musical expectations contribute to increase of arousal and pleasure suggest a possible link with learning based on the relevance of rewarding stimuli and reward-related dopamine circuits ^15^. For example, animal studies show that brain plasticity associated with auditory learning is greater when the information to be learned is rewarded ^61^. Further, pairing a tone with stimulation of dopamine circuits in the brainstem increased the selectivity of responding in auditory neurons tuned to the same tone ^62^. Importantly, dopamine has also been shown to modulate motor learning in humans and animals both directly ^63,64^, and indirectly through monetary reward ^65,66^. Based on this body or work, we test the idea that, by carrying abstract reward, music that is better liked could be associated with greater arousal and learning.

As the motivation to learn a musical excerpt can derive from perceived pleasure, learning can also be motivated by its inherent challenge. Psychological and computational accounts of motivation distinguish extrinsic from intrinsic motivation: whilst the first is based on external reward or pressures outside the individual, the latter is defined as doing an activity for its inherent satisfactions, that have the appeal of aesthetic value or challenge for the individual ^67,68^. It is possible that learning becomes pleasant for actions that lead to decrease of uncertainty and improvement of internal predictive models ^69–71^. Beside this, motivation to learn may be also driven by individual’s feeling of competence in achieving a self-determined goal just for the challenge entailed by the task ^68^. Therefore, individual differences in intrinsic motivation should also be assessed to predict learning.

We tested participants with no-to-little musical training with pupillometry in a listening task, followed by a melody learning task (Figure 1). We composed novel melodies which varied in their structural predictability (from overall predictable to unpredictable melodies). We could formally determine the complexity of the stimuli by means of a variable-order Markov model of melodic expectation (IDyOM)^1^: this model acquires knowledge of musical structure through unsupervised statistical learning and uses this knowledge to estimate the probability of musical notes in a given melody. The expectedness of each note in the melody is expressed in units of information content (IC), where high and low IC values correspond respectively to less and more predictable notes. During the listening task, we recorded the ocular pupil dilation response, and participants were asked to rate how much they liked each melody on a seven-point scale. In the melody learning task, participants learned to play the last four notes of the melodies on a piano-type keyboard (learning phase), and they were tested thereafter (test phase). Importantly, the last four notes did not differ in predictability or motor requirements (See Figure 1), but only in the predictability of the preceding musical context.

**Figure 1.**
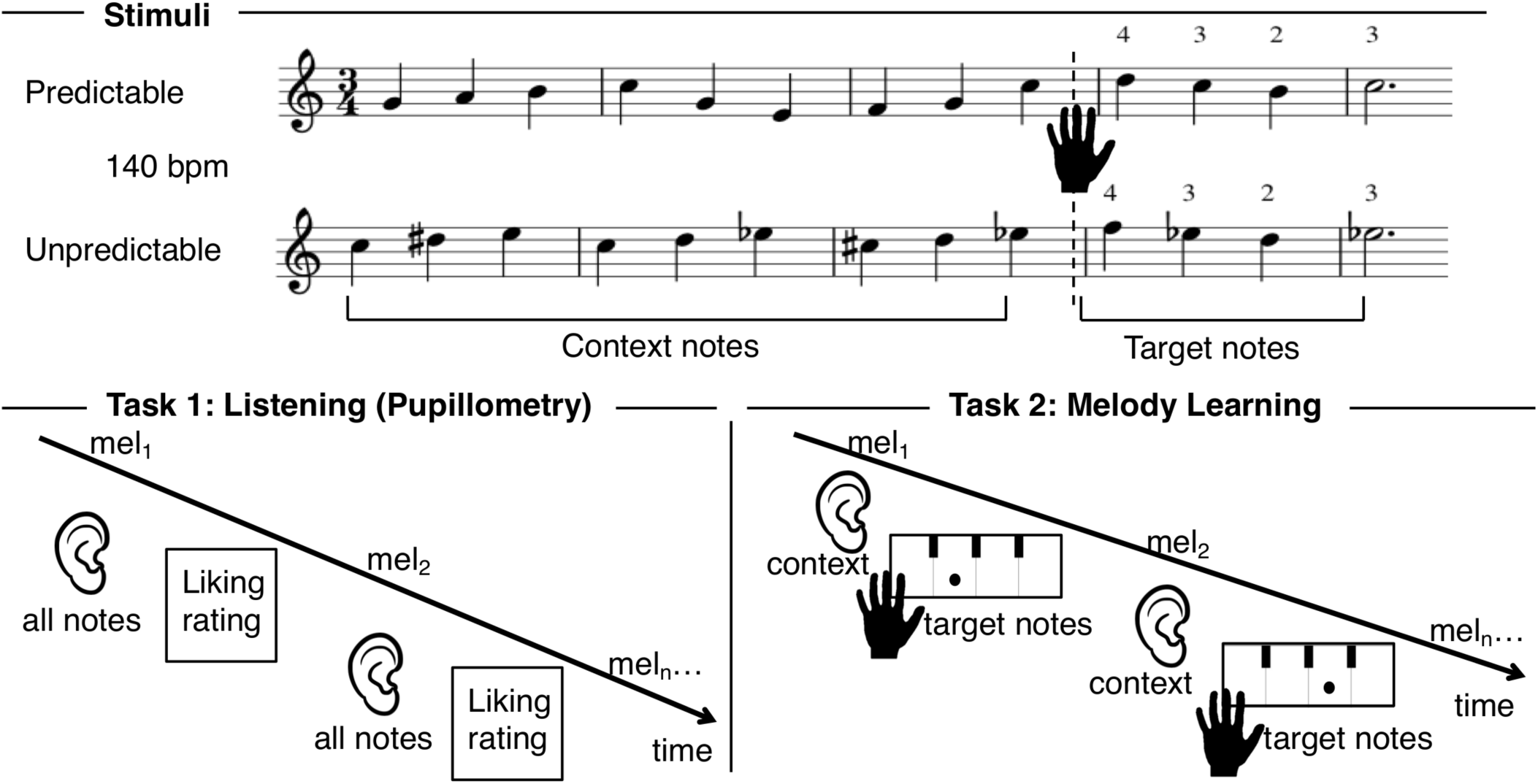
Schematic illustration of the experimental paradigm. **Stimuli.** The musical scores of 2 examples −1 more predictable and 1 less predictable based on the information content (IC) of the context notes. Participants listened to the context notes and then played the target notes as guided by the visual display. The IC of the context notes (before the dotted line) was manipulated to result in predictable (low IC) or unpredictable (high IC) contexts. These were followed by four target ending notes with similar IC between predictable and unpredictable melodies. Fingering for the target notes is indicated by the numbers on the last two bars. Thumb, index, middle and ring finger were assigned to a fixed white key to which different expected sounds were artificially mapped to. This ensured that motor demands for the target ending notes were matched across conditions. **Task 1.** Listening and pupillometry: pupil dilation was measured while participants listened to the entire melody and liking ratings (7-point scale) were collected at the end of each trial. **Task 2.** Melody Learning: participants listened to the first nine context notes of the melody and completed the melody by playing the last four target notes on a midi-keyboard. The notes expected to play were cued by sequential dots drawn onto a keyboard on the screen. Each note occurred at a tempo of 140 bpm, and a metronome sound at 46 bpm (every three notes) guided participants’ pace. Each trial was repeated 5 times during training and 1 time in a final test-phase.

First, we hypothesized that moderately predictable melodies would be better liked, consistent with the inverted-U hypothesis. Further, we predicted that sustained pupil dilation would be greater for melodies that were better liked. For the melody learning task, we expected that predictable musical contexts would result in better motor implementation of target notes, and that better-liked melodies would potentially be better learned even in the unpredictable condition. Because there are large inter-individual differences in music reward sensitivity and intrinsic motivation to perform a task, these characteristics were assessed via questionnaires: Barcelona Music Reward Questionnaire; ^72^ and the Intrinsic Motivation Inventory (with focus on interest/ enjoyment, and perceived competence subscale)^73^.

## 2 Results

### 2.1 Liking ratings of the melodies as function of Predictability

Based on theoretical and empirical work ^47–49^, we expected an inverted U-shape relationship between subjective liking and stimulus complexity. Therefore, a parabola was fitted in a model describing participant’s ratings (scaled by subject) as a function of IC of melodies (averaged across notes). Figure 2 shows the quadratic relationship between melody IC and participants’ liking ratings (*χ*^*2*^(1) = 4.513, *p* = .033), indicating that liking was higher for moderately predictable melodies, but it decreases when melodies become more complex (high IC). Possibly because of a relative narrow range of IC across melodies, a linear model could also fit the relationship between liking and IC (*χ*^*2*^(1) = 4.174, *p* = .041), which yielded similar model fit to the quadratic term (ΔAIC = .6).

**Figure 2.**
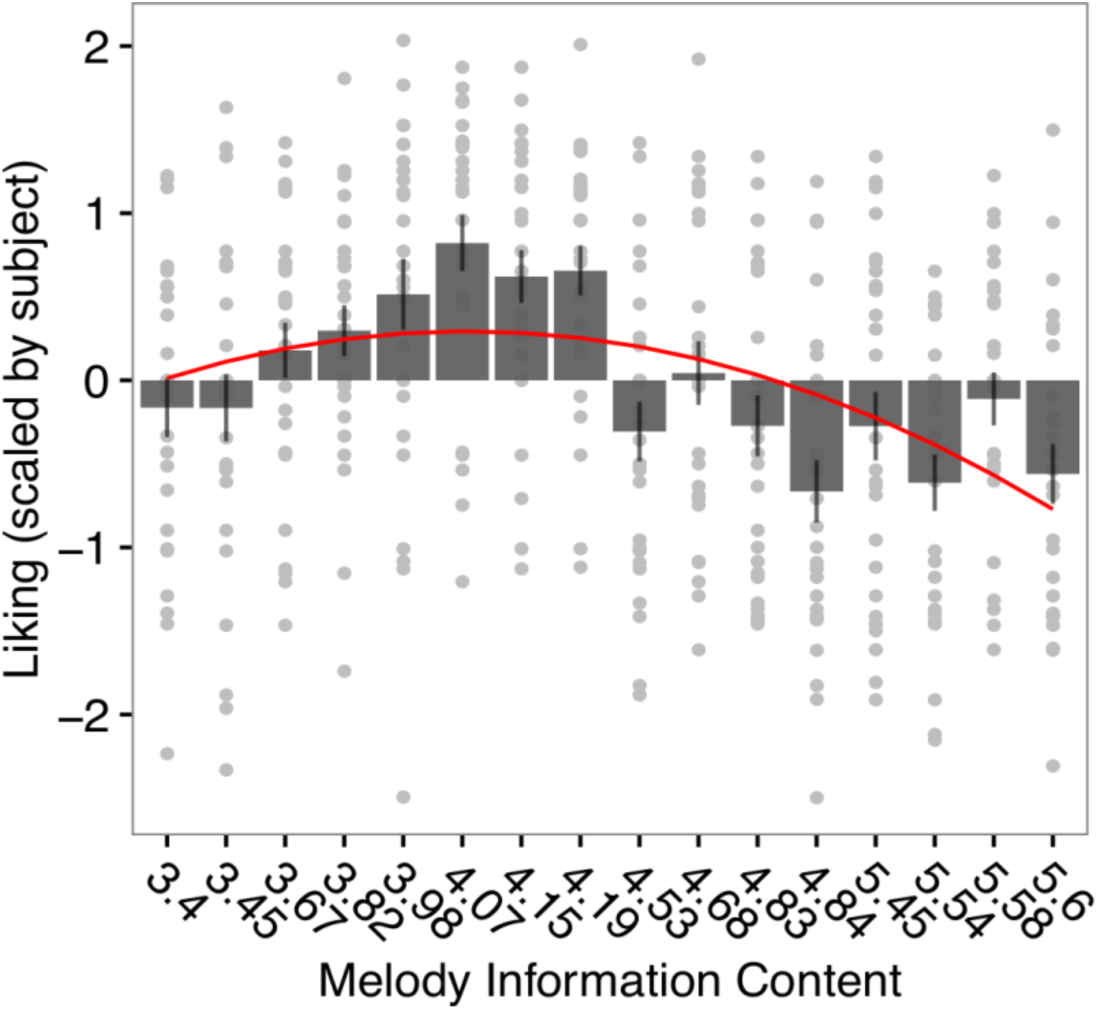
Inverted-U relationship between mean information content (IC) of each melody (ordered by increasing mean IC values on the x axis) and subjective liking ratings (scaled by subjects). Each point represents individual ratings for each melody. Error bars represent 1 s.e.m. of ratings for each melody.

### 2.2 Pupil diameter during listening

During listening, participants provided liking ratings to each of the melodies whilst pupil dilation was measured. We analyzed the effects of predictability (Predictable/Unpredictable) and liking (High/Medium/Low) and their interaction on participants’ pupil size change over 13 time-bins, corresponding to the 13 notes in the melodies (Figure 3). We found an effect of predictability by time bin on the sustained pupil response (*χ*^*2*^(1) = 5.291, *p* = .021). Although the differential increase over time did not reach statistical significance after multiple comparisons, post hoc paired t-test showed overall greater pupil dilation for predictable compared with unpredictable melodies [P-U: *t*(1,22)= 3.137, *p* = .005]. Moreover, we found an interaction of liking and time bin (*χ*^*2*^(2) = 29.257, *p* < .001): this indicated that high-liked melodies induced greater dilation than medium-liked (high-medium: *b* = 5.942, SE = 1.458, *p* < .001), or than low-liked melodies (high - low: *b* = 3.928, SE = 1.469, *p* = .021). A three-way interaction of predictability, liking and time bin (*χ*^*2*^(2) = 6.152, *p* = .046) showed that pupil dilation for the high-liked condition increased more for predictable than unpredictable melodies (P - U: *b* = 4.846, SE = 2.107, *p* = .021). No predictability effect was indeed found in the low-liked (P - U: *b* = -1.113, SE = 2.05, *p* = .587), or medium-liked condition (P - U: *b* = -1.783, SE = 2.19, *p* = .377). These results suggest that the increase of pupil dilation as a function of liking was greater when the music was predictable.

**Figure 3.**
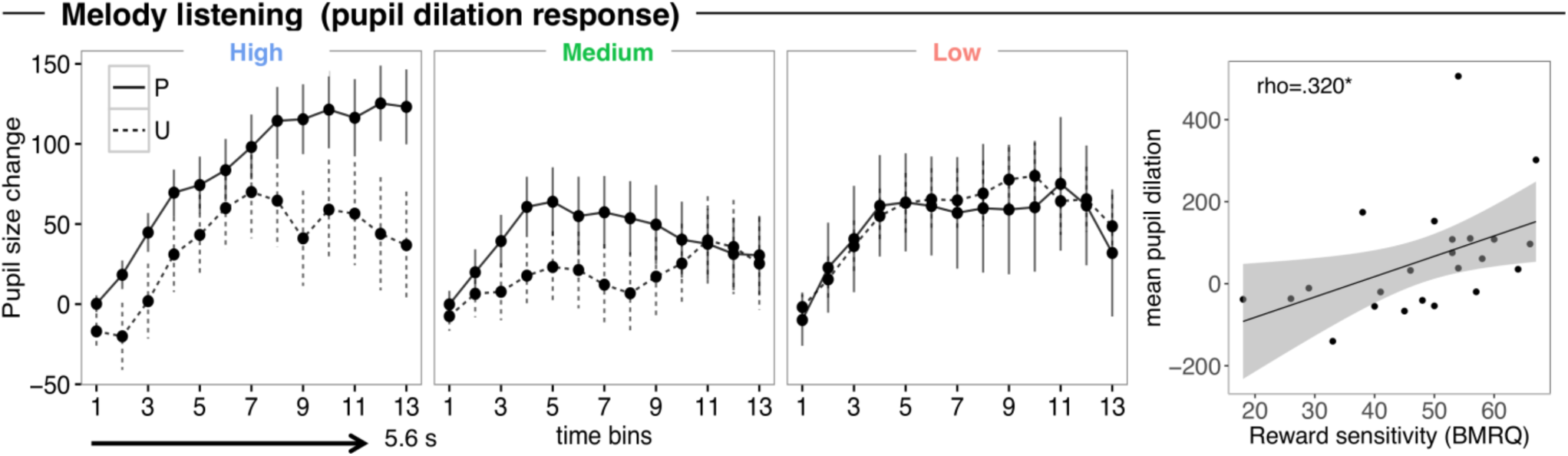
Pupil size change across 13 time-bins (for each note) for predictable (P) and unpredictable (U) melodies across different degrees of liking (high, medium and low). Error bars represent 1 s.e.m. of all trials. (right panel) Scatter Plot showing the mean pupil dilation as a function of reward sensitivity score. Each data point represents an individual participant. The diagonal indicates the line of best fit.

Finally, individual scores of music reward sensitivity positively predicted pupil size change across all trials regardless of condition (rho=.320, *p* = .005) (Figure 3; right panel), showing that individuals who are more sensitive to musical reward have a greater physiological response.

### 2.3 Motor performance

In the motor task participants listened to the first nine *context* notes of each melody through headphones and then played the last four *target* notes on a piano-type keyboard. Accuracy and asynchrony were entered in separate mixed effects regressions for the training and the following test phase.

In the training phase, we found increase of *accuracy* across repetition trials (*χ*^*2*^(3) = 22.498, *p* < .001), and no other effect involving Predictability or Liking (*p*s > .111). For *asynchrony* (Figure 4), there was a main effect of Predictability (*χ*^*2*^(1) = 11.677, *p* < .001), and an interaction with repetition trial (*χ*^*2*^(1) = 5.758, *p* = .016), such that predictable trials were better executed, and that this advantage were greater in the early trials (P – U across repetition trials: *b* = 5.883, SE = 2.753, *p* = .033). An interaction between Liking and Predictability (*χ*^*2*^(2) = 9.092, *p* = .01) indicated that these effect of predictability mainly regarded the medium-liked melodies (P - U: *b* = -27.872, SE = 11.454, *p* = .015), but not the high-liked (P - U: *b* = -5.675, SE = 11.659, *p* = .131), and low-liked ones (P - U: *b* = -17.484, SE = 11.574, *p* = .131). The poorer performance for medium-liked melodies in the unpredictable condition suggests that the unpredictability of the stimulus is compensated when strong (positive or negative) rather than mild (medium) emotional responses are at play. We indeed found a significant main effect of Liking (*χ*^*2*^(2) = 16.755, *p* < .001), and an interaction with repetition trial (*χ*^*2*^(2) = 21.263, *p* < .001) such that learning was better for the high-liked and the low-liked melodies when compared with the medium-like ones (across repetition trials high-medium: *b* = -12.718, SE = 3.401, *p* < .001; low-medium: *b* = -7.998, SE = 3.313, *p* = .016; high-low: *b* = -4.720, SE = 3.398, *p* = .165).

**Figure 4.**
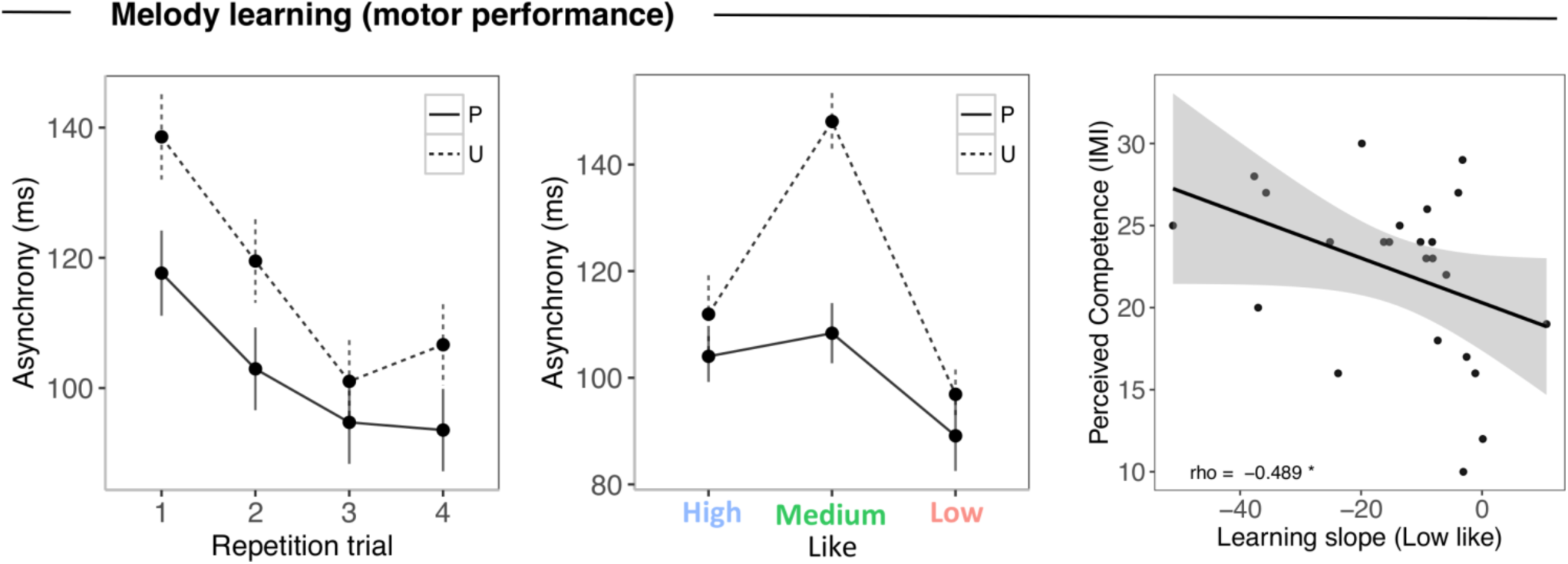
Training phase. Average asynchrony for 4 repetition trials for the keystroke of target notes in predictable (P) and unpredictable (U) melodies (left panel). Average asynchrony for predictable and unpredictable melodies across different degrees of liking (high, medium and low) (middle panel). Error bars represent 1 s.e.m. of all trials. (right panel) Scatter plot showing the correlation between intrinsic motivation score of perceived competence and learning regression slope for Low Like melodies. Each data point represents an individual participant. The diagonal indicates the line of best fit.

Taken together, these findings indicate that learning of the target-notes was greater after predictable than unpredictable contexts in presence of mild emotional responses. However, the disadvantage due to the unpredictable contexts was minimized for high- and low-liked melodies, which showed similar learning regardless of predictability.

We further investigated the role of other factors than stimulus-induced affective responses in learning, as inter-individual differences associated with intrinsic motivation. Specifically, we tested the relationship of the general enjoyment and task-perceived competence IMI scales, with the learning slopes – measured as reduction of asynchrony across repetition trials– in the high-like and low-like conditions. To estimate the individual performance improvement across repetition trials as a function of liking, we ran mixed model testing for only the effect of liking (High/Medium/Low) across repetition trials, and extracted residual individual slopes for the low-/high-liked condition, after adjustment for the effects of the performance change in the medium like condition at the intercept. More negative slopes (faster learning) associated with low-liked condition were predicted by higher scores of individuals’ perceived competence in succeeding the task (Fig. 4 right panel, rho = -.488, *p* =.015), and there was a trend with the general enjoyment scale (rho = .393, p =.085). The learning slope for high-liked melodies did not correlate with perceived-competence scores (rho = -.224, p =.292), nor with general enjoyment scale (rho = .147, p =.535). Thus, the learning effect observed in low-liked melodies was particularly driven by participants with high task perceived competence, whilst liked music was learned regardless. This suggests that perceived competence helped the participants overcome low liking to learn the melodies, but it didn’t play a role in highly liked stimuli as participants were already motivated by the pleasure carried by music.

Furthermore, we tested if learning highly liked stimuli was predicted by pupil response to those stimuli. The correlation between the mean amplitude of pupil diameter in high-like trials and the learning slope of high-like condition did not yield significant results (rho = .044, p =.84). This may be due to the fact that the range of liking values was relatively limited.

In the test phase, we didn’t observe any significant effect of liking nor predictability on accuracy and asynchrony (all *p*s >.389). This non-finding in the test phase may be explained by the fact that only one trial per melody was not sufficient for seeing an effect. Also, the increased number of errors committed in the test phase compared with the last repetition trial of the training phase (pitch and timing errors combined, mean averaged across participants 10.74 ± 26.60 %, in the last repetition trial; 35.51 ± 24.04%, in the test phase; t (25) = -5.940, *p* < .001, Cohen’s *d*=-2.376), suggests that more repetitions may be desirable for stabilization of learning.

## 3 Discussion

The present study investigated the contribution of predictability and liking on arousal and learning in non-musicians. First, we found an inverted U-shaped relationship between music complexity and liking, showing that moderately predictable melodies were more liked than highly predictable and unpredictable melodies. Further, we showed a synergistic effect of predictability and hedonic response to music on arousal, as reflected by sustained pupil dilation. We also observed that pupil dilation was overall greater in individuals with higher sensitivity to musical reward, suggesting that it is a good marker of responsiveness to music. With regard to melody learning, performance was facilitated for predictable compared to unpredictable melodies in medium-liked music, indicating that musical expectations can facilitate auditory-motor predictions and movement preparation, even in non-musicians. This effect of predictability was overshadowed by musical reward as liked melodies were better learned, even when unpredictable. Finally, we found that not-liked melodies were also learned and that this effect was correlated with individuals’ task perceived competence, suggesting that, beyond the musical reward, other factors carrying reward — such as individual’s task-related intrinsic motivation— contribute to learning.

Our results link quantitative measures of stimulus complexity (as music predictability) with liking response of listeners as an inverted-U-shaped function, whereby moderately predictable melodies were more liked than highly predictable and unpredictable melodies. Importantly, listeners were unfamiliar with the stimuli, and complexity was objectively characterized by the IDyOM model which has been shown to optimally predict subjective perceptual expectations ^74^, and perceived complexity of musical structure ^75^. The inverted-U model was first proposed by Berlyne (1971) to reflect a general relationship between aesthetic appreciation and structural complexity in art. But, it has also been shown to be a general property of complex stimuli including visual shapes ^76^, music and rhythm ^44,49^. In line with a predictive account, liking may derive from an intrinsic reward which occurs whenever an internal predictive model improves by decreasing uncertainty ^69,71^. Because the potential for decreasing uncertainty is maximal when music is moderately complex, so it should be for the liking. The inverted-U model has received empirical support in some music experiments ^47,48,50^, but not in others ^77,78^, possibly because it is often difficult to generate ecologically valid stimuli that cover the full range of complexity, or because other psychological mechanisms triggered by the stimulus such as familiarity, imagery, memory or associations ^1,79^ interact with expectation-based emotions.

The effect of liking on subject’s arousal is in line with previous literature ^14^, in that pupil response during listening was greater for liked melodies. Importantly, we extend this finding by showing that pupil dilation increased for high-liked melodies, but only when they were predictable. These results are novel because they address for the first time both the effect of subjective pleasure and musical complexity on sustained pupil response over relatively long stimuli. They are compatible with the interpretation that sustained pupil dilation is modulated by both changes in attentional engagement due to stimulus structure tracking ^80,81^, and to subjective affective evaluation or reward ^14,82^. In support of this interpretation, electrophysiological evidence has established a link between pupillary response and norepinephrine activity in the nucleus locus coeruleus ^83^ that has synergistic connections with subcortical dopaminergic nuclei involved in reward, and prefrontal areas involved in stimulus evaluation processes ^55,56^. Predictable melodies may thus result in greater attentional engagement – enhanced noradrenergic activity – as they conform to listeners’ prior expectations and allow them to form precise predictions about the incoming stimulus ^1^. Such effect of predictability may explain effects of greater sensitivity and memory of music from one’s own culture, or of simple more than complex excerpts ^31,32^. Conversely, unpredictable melodies may down-weigh predictions from a model that does not match the incoming stimulus, resulting in attentional disengagement, and lower pupil response. The interaction effect of liking and melody predictability on pupil dilation suggests that positive evaluative processes build on successful tracking of the stimulus structure. One proposed mechanism is based on a hypothesized feedforward loop between forebrain regions associated with reward evaluation and the concerted action of the noradrenergic and dopaminergic systems ^55^. As a positive subjective evaluation is formed throughout the melody, succeeding valid predictions gain greater reward value through dopamine-mediated response, which in turn boosts norepinephrine-mediated attention. Future investigations combining pupillometry and brain imaging are necessary to identify this circuit, and the proposed dynamic interaction during response to music.

Learning of the target notes was facilitated for predictable compared with unpredictable melodic contexts in medium-liked music, demonstrating that musical structure promotes predictions and motor encoding in naïve performers. Importantly, better learning cannot be explained by differences in predictability or motor complexity intrinsic to the target notes, because these were similar across all melodies. They only differed because they were embedded in contexts that allowed better or worse prediction of the most likely continuation of the melody. Further, learning effects were observed for temporal accuracy of the movements, not note accuracy – which was high in all conditions. This is important because the stimuli varied in melodic expectations, but not in timing. Thus, better temporal accuracy for more predictable melodic contexts indicates that musical structure promotes motor prediction and planning by heightening the precisions of the movements. This result is in line with the notion of ‘active inference’ ^22,24^: by relying on models of the environment with a high level of precision in predictable contexts, the brain can select a narrower set of information to predict the future sensorimotor state and to reduce uncertainty ^84,85^. Given that perception and action are intertwined, perceptual and motor networks may also interact during the generation of predictions about the most likely next state ^26^. Accordingly, in music models of musical structure inform the sensory system to anticipate the most predictable sound ^86^, and they also drive the motor systems to facilitate the movement required to produce it ^37,39,42,87^. Moreover, there is recent evidence that non-musicians rapidly form sensorimotor representations of anticipated events after short-term motor training ^36,88^. Our results suggest that even in naïve performers, predictions based on experience in the auditory modality affect predictions in the motor domain, irrespective of previous training linking sounds to actions. A possible underlying mechanism may be the rapid formation of sensorimotor associations at the first attempts of execution, which result in facilitated performance in the following repetition trials^89^. They may also be based on long-term priors –the so-called SMARC effect (Spatial Musical Association of Response Codes) – which shows that even for individuals without training, higher pitches facilitate upward or rightward responses, and low pitches facilitate downward or leftward responses ^90^. Alternatively, in line with the view that sensory and motor systems act as independent “emulators” of upcoming events ^91^, predictive models in the motor domain may be independently generated based on existing models of music built through auditory perception.

Liking a melody reduced the disadvantage in performance due to the unpredictable contexts, suggesting that music-induced hedonic response promotes learning by overshadowing the effect of predictability. A possible underlying mechanisms may be an interaction between dopamine-mediated reward induced by music ^71^ and dopamine-mediated learning mechanisms ^15^. This is consistent with work reporting enhanced motor learning and retention in presence of external incentives, such as monetary reward ^65,66^. In line with the idea the reward value of music may act as a reinforcement signal for learning ^51^, our results foster the link between reward and motor learning in a more complex task and for an abstract stimulus-related incentive.

The motor learning benefit associated with preferred music may be indirectly linked with general greater attention, as reflected by increase of pupil in liked melodies. The well-known interaction between the noradrenergic system – underlying pupil dilation – and the dopamine system – associated with reward – ^55,92^, suggest that the concerted action of these two systems may mediate the beneficial effect of music reward on memory and motor learning. We did not find evidence to relate motor learning and pupil response to liking ratings, probably because of the limited range of response elicited by the stimuli used here. Previous studies using stimuli that induce musical chills have shown that they induce greater pupil dilation ^14^, and are also better remembered ^57^, consistent with the key role of the reward system in stimulus encoding ^93^. Future studies, likely in trained musicians, could use stimuli which induce more intense pleasure to examine their effect on learning.

Non-liked melodies were also learned similarly to liked melodies, irrespective of music predictability. The learning of non-liked melodies was driven by participants with higher task-achievement motivation (perceived-competence scale), as opposed to a general learning effect of high-liked melodies. This is in line with the definition of competence where the achieving process, rather than goal being achieved, is central ^67^. Thus, our results suggest that when the music is not rewarding per se, people with greater general task-related motivation succeed better in learning it.

Predictability and liking are inherently linked in music. Their intertwined effect was evident in the pupillary response, which was enhanced both by musical expectations and subjective music reward. However, what are their contributions to learning when assessed separately? We observed that when operationalized as information content, effects of predictability on learning can be over-shadowed by effects of liking or of intrinsic motivation. One implication of this is that liking in music should neither be reduced to “mere” liking – as it can drive learning, thus acting “as” a reinforcer, nor to mere predictability because unpredictable melodies were learned equally well when liked. This may also in part be due to the fact that subjective liking plausibly involves many dimensions beyond predictability, such as familiarity, imagery, memory or idiosyncratic associations ^1,79^. These results reinforce the view that emotional and motivational factors have powerful impact on learning not only for cognitive tasks, but also for procedural and motor-skill learning ^94^. In conclusion, this study provides an important first step in understanding how motor learning benefits from the contributions of implicit musical expectations and the derived emotional response. Future research in this direction may shed light on their additional benefits on rehabilitation in clinical populations.

## 4 Material and methods

### 4.1 Participants

Twenty-seven individuals with no previous piano training took part in the study (18 female; Age: *M*= 24.74 ± 5.03). Participants had on average less than one year (*M* = 0.8 years + 1.5) of formal music training, which did not take place in the last 10 years (note that one participant with 16 years of training in another instrument was excluded from the analysis). All participants were neurologically normal, were not taking any medication that could affect motor performance, and had normal hearing, and normal or corrected to normal visual acuity. All participants were naïve with regard to the purpose of the study and provided written informed consent. The Concordia University Human Research Ethics committee approved the study (30007730) and conducted adhering to the Canadian Tri-council Policy on ethical conduct for research involving humans ^95^.

### 4.2 Stimuli and Procedure

#### 4.2.1 Stimuli

Sixteen different melodies of 13 notes were newly composed specifically for the experiment according to the rules of classical Western tonal music. All of them began on the first beat, and were notated with a time signature of 3/4 that was thought to be easier to count along. In order to focus specifically on pitch expectations, each note had the same duration and equivalent inter-onset interval of 428 ms (140 bpm). Melodies were created with MuseScore program (version 2.0.2) and synthesized with a piano sound (generated using Ableton Live 8) with the same loudness for all notes and melodies. Each melody had a total duration of 6.4 s.

The predictability of each melody was objectively defined using the information dynamics of music model, IDyOM ^19^ and based on the average information content values of each note. This model is trained through a process of unsupervised learning on a large training set of 903 Western tonal melodies. IDyOM first analyses the statistical structure of the training set, represented as sequences of pitch and note’s scale degree relative to the key of the melody. In a new sequence, it then estimates the probability of each note, based on a combination of the training set’s statistics and those of the sequence at hand, which it learns dynamically. The output is a note-by-note measure of information content (IC, the negative logarithm, to the base 2, of the probability of an event occurring), which IDyOM uses instead of raw probability for greater numerical stability and a meaningful information-theoretic interpretation in terms of redundancy and compression. We manipulated the mean predictability/IC of each melody by varying the number of out-of-key notes over the first 9 *context* notes – whereby an out-of-key note results in high IC (Figure 5A). The predictability/IC of the four *target* notes designed to be similar across melodies (Figure 5B). An ANOVA with factors Predictability (P/U) and Note type (Context/Target Notes) on the IC of each note yielded a significant interaction of Predictability and Note type [F_(1,14)_ = 11.60, *p* = .004, η p^2^ = .45], indicating that IC for Context but not Target notes differed significantly between predictable and unpredictable melodies [Main effect of predictability on context notes only: F_(1,14)_ = 52.88, *p* < 0.001, η_p_^2^ = .79; Main effect of predictability on target notes only: F_(1,14)_ = 2.50, *p* = .136, η_p_^2^ = .15]. Based on the IC measure the 16 melodies were ranked from high to low probability (*M*= 4.48 ± 1.56, range = 1.7-9.1), and then they were divided into Predictable or Unpredictable based on the median split (Figure 5A).

**Figure 5.**
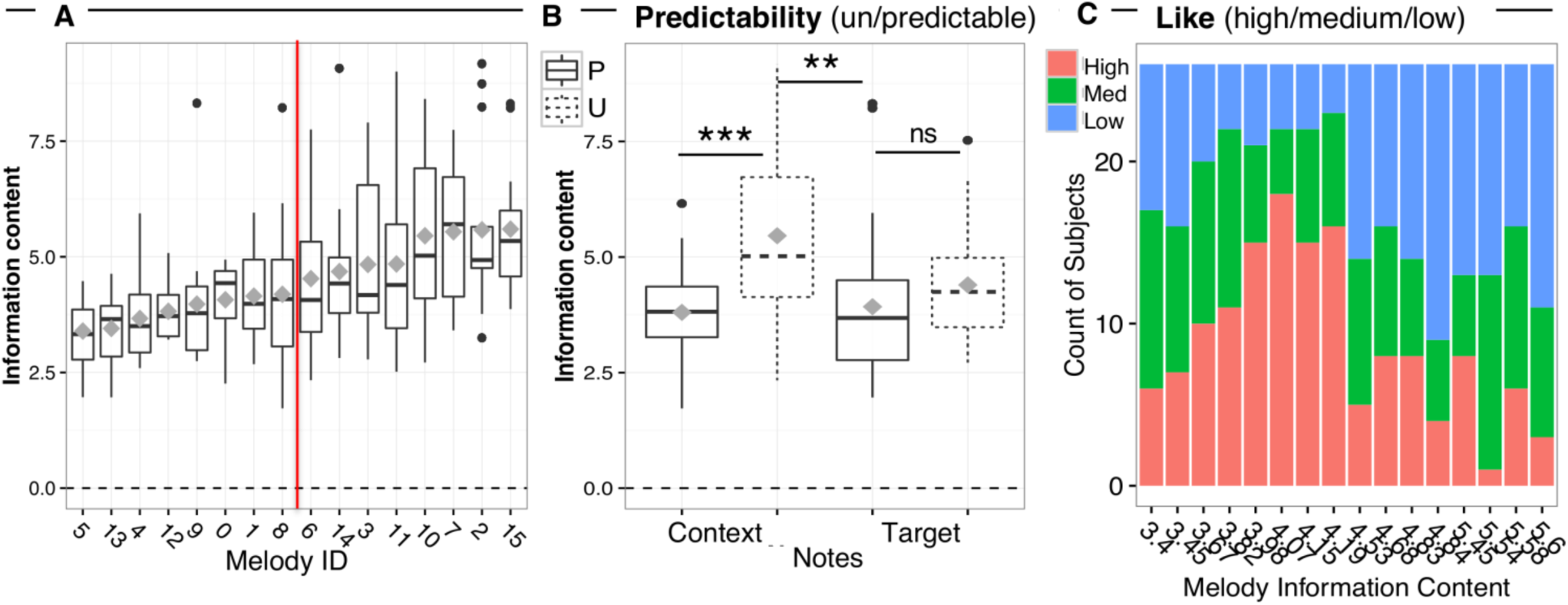
Stimuli. Median information content A) for each melody, B) for context and target notes of predictable (P) vs. unpredictable (U) melodies. Box = 25th and 75th percentile; bars = min and max values. Grey squares = mean information content across notes. The red vertical line indicates the median split of melodies based on their mean information content C) Stack bar chart with the counts of high, medium and low liking ratings assigned by subjects to each melody (on the x axis ordered by increasing mean IC values).

#### 4.2.2 Liking ratings and pupillometry

Participants listened to all 16 melodies, one at a time, and rated how much they liked them using a 7-point Likert scale (1 being not all and 7 being very much) at the end of each melody. Liking ratings for each melody were then scaled by subject, and ranked as high, medium and low (Figure 5C).

Pupil dilation was measured during Listening using the EyeLink 1000 head-supported infrared optical eye-tracking system (running host software ver. 4.56, SR Research, Ottawa, ON, Canada) in binocular 1000Hz sampling configuration, connected to an Apple iMac (Mac OS X 10.12). The EyeLink system was used in the Pupil-Corneal Reflection tracking mode. Participants were seated in a comfortable chair with their head stabilized in a chin and forehead rest, facing the computer monitor (View sonic G225fb 21” CRT, 1024 × 768pixel resolution, 100 Hz refresh rate, linear gamma correction for luminance with mean luminance = 60 cd/m^2^) at a distance of 70 cm, in a quiet, moderately lit room (40 cd/m^2^). To calibrate the eye-tracker, a circular target (1 degree of angle) appeared in random order at one of 6-points on the screen (HV6, in the default Eyelink screen locations), followed by a separate validation. Calibration and validation were repeated until the average error across all calibration targets was below 0.5 degrees of visual angle, and the maximum error at any one calibration point was below 1 degree of visual angle. The order of presentation of the melodies was randomized across participants. Two seconds of baseline pupil data was acquired before and after each melody was played. After each melody, two seconds were given for the liking rating, which was followed by a 2-seconds blank grey screen before the next trial. Participants were instructed to continuously fixate on a cross at the center of the screen (size: 9.4cm x 9.4cm, corresponding to a 4.5° visual angle at a viewing distance of 70 cm; RGB: 75,75,75; background grey color, RGB: 150,150,150), not to move their heads for the duration of the eye tracking component of the experiment, and to avoid blinking while the melody was playing. The total duration of the Listening task was approximately 20 minutes.

#### 4.2.3 Melody Learning

In this task participants listened to the first nine *context* notes of each melody through headphones and then played the last four *target* notes on a piano-type keyboard using the four fingers of the right hand: thumb, index, middle and ring finger which were assigned to a fixed white key to which all notes were mapped. This ensured that motor demands for the notes to be played were matched across conditions. Target notes were cued using visual display representing the keyboard (see Figure 1) where a dot appeared sequentially (IOI 428 ms) to indicate which key to play. Participants heard the notes they produced through the headphones. To facilitate accurate playback timing, melodies were accompanied by a metronome beat at the beginning of every bar. The fourth metronome beat cued participants to begin playing back the target notes. The notes played and their timing were recorded from the keyboard and used to score accuracy and synchronization. The 16 melodies were presented in a randomized order, and each melody was repeated 5 times with an ISI of 1 second. At the end of the Learning task, participants performed a final recall block where each of the sixteen melodies was played back once in a random order.

Before training, participants were familiarized with the playback task in a brief practice block of four trials in which they had to count 3 metronome beats (corresponding to 3 bars), and then perform 4 keypresses cued by the visual display. The familiarization trials contained all finger transitions that were to be encountered in the Melody Learning task. No auditory feedback was provided. Presentation of the melodies and recording of the responses was controlled custom-written Python software running on a PC Linux computer.

At the end of the experiment, each participant’s music reward sensitivity (i.e. how important music is in his/her life) was assessed via the Barcelona Music Reward Questionnaire ^72^. Participant’s intrinsic motivation to perform the task was assessed via the standard, 22-items version of Intrinsic Motivation Inventory ^73^. The entire experiment lasted approximately 60 minutes.

### 4.3 Data analysis

#### 4.3.1 Analysis of pupil diameter

Pupil diameter was measured in arbitrary units. Blinks were identified by identifying in each trial samples without data due to blinks, removing 100 ms before and after the edges of the non-data points to make sure that all artefacts of the pupil size algorithm were removed. For each blink, four equally spaced time points (t2 = blink onset; t3 = blink offset; t1=t2-t3+t2; t4=t3-t2+t3) were interpolated by using a cubic-spline fit and the original signal was replaced by the cubic spline, leaving the signal unchanged except for the blink period. Random sample artefacts were removed using a median Hampel filter (from EEGlab software, version 14.1), after which data were smoothed using a Savitzky-Golay Filter over an 11-ms timeframe to remove the high-frequency noise in the pupil without time-delaying the pupil signal. Then, each trial was baseline-corrected against the median pupil size in the 400 ms before the onset of the melody, and then divided in 13 bins corresponding to the onset of each note. Trials during which participants blinked for more than 15% of the total trial duration were excluded (two participants’ datasets, and a mean of 0.92 ± 2.53 for the rest of participants). Baseline-corrected pupil size change was then analyzed by using linear mixed-effects regressions testing for the effects of predictability (Predictable/Unpredictable), Liking (High/Medium/Low), time-bins (1-13) and their full interaction.

#### 4.3.2 Analysis of motor performance

Participants’ performances were examined off-line to evaluate key errors (MIDI note number) and response times for the four keystrokes relative to the four last target notes of each melody. Trials were considered invalid and excluded from the analyses if participants pressed more or fewer than 4 notes per trial. On these data, two indexes of performance were computed: *trial accuracy* (i) was quantified by counting the total number of errors. These were defined either by an incorrect keystroke (key identity error), or by an absolute response time larger than 428 ms (timing error; for values outside this range, keystrokes occurred within the range of the note preceding or following the one with which it was supposed to be synchronized). *Asynchrony* (ii) was quantified only on correct trials by measuring the time difference between the actual keystroke of a note and the expected onset of that given note.

Statistical analyses were performed separately for the two performance indexes (i.e., trial accuracy and asynchrony) and for the training (where each melody was performed for 5 consecutive times) and the test phase (where each melody was played only one time). We used mixed effects regression analyses testing for the effects of predictability (Predictable/Unpredictable), liking (High/Medium/Low), repetition trial (1-4), keystroke (1-4), and the full interaction between liking, predictability and repetition trial. Keystroke was introduced as an effect of no interest to account for known motor execution differences between initiation (first key press) and completion (following 3 keypresses) of sequential movements ^96^. Keystrokes for the first repetition trial were initially analyzed, but then excluded because of too many invalid trials (*M*= 55.04 ± 26.60 % of invalid trials across participants). The analysis on the test phase looked for learning stabilization, and estimated the effects of predictability (Predictable/Unpredictable) and liking (High/Medium/Low).

#### 4.3.3 Statistical analysis

All data analyses were conducted in MATLAB R2015b (Mathworks, Natick, MA, USA), except for the linear mixed model analysis that was implemented in R environment Version 0.99.320 using the ‘lmer’ function from package lme4 to build the models ^97^ and the ANOVA function from package car to obtain significance tests ^98^. In contrast to a more traditional approach with data aggregation and repeated-measures ANOVA analysis, linear mixed effects regression allows controlling for the variance associated with random factors without data aggregation (see ^99^). By using random effects for subjects and stimuli item, we controlled for the influence of different mean responses associated with these variables. Moreover, we also included by-participant random slopes for the effects of interest (predictability, liking and their interaction), which accounted also for differences in how predictability and liking affected participants’ responses (random slopes). Contrasts were carried out using the ‘emmeans’ package in R^100^. We report unstandardized effect sizes (unstandardized regression coefficients, indicated as ‘b’ for the statistical tests) which is in line with general recommendations of how to report effect sizes in linear mixed models ^101^. Significance of the fixed effects of these models were evaluated with the Satterthwaite approximation ^102^, and *p* values were adjusted for multiple comparisons using the multivariate t method.

## 6 Author Contributions

**RB** conceived performed and analyzed the experiments; wrote manuscript.

**BG** provided expertise and feedback; commented on manuscript draft.

**AJ** provided expertise and feedback on pupillometry and analysis; commented on manuscript draft.

**VP** supervised and administered the project; secured the funding; conceived the experiments; wrote the manuscript.

## 8 Acknowledgments

We are grateful to Joe Thibodeau for his assistance in developing the hardware and software interfaces for this experiment. We acknowledge the contribution of Soraya Lalou in collecting the data. We also thank Prof. Robert Zatorre and Dr. Gabriele Chierchia for inspiring discussions, and three anonymous referees for very constructive suggestions.

## 9 Data Availability Statement

The datasets and the stimuli for this study can be found in the OSF repository (link: https://osf.io/x42sz/?view_only=e75f0dd5b6964c3cb39f603095141885).

## 10 Informed Consent Statement

All participants provided written informed consent. The Concordia University Human Research Ethics committee approved the study (30007730) and conducted adhering to the Canadian Tri-council Policy on ethical conduct for research involving humans

